# Engineered protein-small molecule conjugates empower selective enzyme inhibition

**DOI:** 10.1101/2021.05.05.442679

**Authors:** Andrew K. Lewis, Abbigael Harthorn, Sadie M. Johnson, Roy R. Lobb, Benjamin J. Hackel

## Abstract

Specific, potent ligands drive precision medicine and fundamental biology. Proteins, peptides, and small molecules constitute effective ligand classes. Yet greater molecular diversity would aid the pursuit of ligands to elicit precise biological activity against challenging targets. We demonstrate a platform to discover protein-small molecule (PriSM) hybrids to combine unique pharmacophore activities and shapes with constrained, efficiently engineerable proteins. A fibronectin protein library was yeast displayed with a single cysteine coupled to acetazolamide via a maleimide-poly(ethylene glycol) linker. Magnetic and flow cytometric sorts enriched specific binders to carbonic anhydrase isoforms. Isolated PriSMs exhibited potent, specific inhibition of carbonic anhydrase isoforms with superior efficacy to acetazolamide or protein alone including an 80-fold specificity increase and 9-fold potency gain. PriSMs were engineered with multiple linker lengths, protein conjugation sites, and sequences against two different isoforms, which reveals platform flexibility and impacts of molecular designs. PriSMs advance the molecular diversity of efficiently engineerable ligands.

## Introduction

Molecular targeting ligands empower precision medicine and fundamental biological research.^1,2^ A multitude of platforms have effectively empowered ligand discovery including computational design^3^ and combinatorial library screening via phenotype-genotype linked display methods^4^ for proteins as well as structure-guided screening^5^ and fragment-based drug discovery^6^ for small molecules. Proteins have large surface area to drive affinity^7^ and specificity^8^, structural organization to limit entropic cost upon binding^9^, and are ribosomally encoded to permit high-throughput genotype-phenotype selections. Small molecules provide smaller size that is more amenable to target concave epitopes, broader chemical diversity, and different opportunities for shape complementarity. These complementary properties, as well as different advances in discovery platforms, have led both molecular classes to yield successful ligands for an array of applications.

Nevertheless, the ability to target select epitopes and specifically elicit desired biological activities remains challenging in some systems, including enzymes and highly homologous targets, which has inspired an array of advanced molecular designs and engineering strategies. Extended antibody complementarity-determining regions^10^ and alternative scaffold topologies^11^ have facilitated cavity targeting. Hybrid molecules with small molecule pharmacophores conjugated to peptides have been engineered to increase specificity on kinases^12–14^ and to increase affinity or potency against lectins^15^, enzymes^16,17^, and microbes^18^.

A key challenge in these latter systems is to efficiently engineer the peptide and conjugated small molecule to form a synergistic binding interface. Peptide-pharmacophore hybrids have been identified via design^19^ and discovered from combinatorial libraries^14,15,17,20,21^. In the current work, we evaluated the ability to merge the benefits of small molecule pharmacophores with folded protein scaffolds, which provide greater surface area, more diverse topology, and reduced entropic costs relative to previous peptide-based hybrids. We evaluated the implementation of protein-pharmacophore hybrid engineering within the context of yeast surface display, which enables quantitative flow cytometric selection for affinity^22^ and specificity^23^; multivalent display for avid selection^4^ of moderate affinities to begin the discovery and evolutionary process, and eukaryotic processing for effective folding. Previous work by Van Deventer and colleagues has demonstrated the ability to conjugate small molecules to non-canonical amino acids in yeast-displayed proteins^24,25^. In the current work, we used the free thiol residue of a single cysteine moiety to enable focused conjugation of a small molecule pharmacophore.

The approach can be considered as an analog of fragment-based drug discovery^26^ in which the pharmacophore extends the binding capabilities of an engineered protein. As another perspective, an integrated binding site comprising protein-target and small molecule-target interfaces builds on the success of heterobivalent AND gates^27,28^, which avidly couple two interactions of moderate intrinsic affinity to achieve high specificity and strong macroscopic affinity. In a third perspective, the conjugated small molecule can be viewed as a non-canonical amino acid to expand the natural repertoire^29^ without the need to engineer translational machinery.

As an initial test system for the <u>pr</u>otein-<u>s</u>mall <u>m</u>olecule (PriSM) hybrid engineering platform, we aimed to selectively inhibit enzymes of both fundamental and clinical importance with off-target homologs. We chose carbonic anhydrases (CAs), a family of zinc metalloenzymes that catalyze interconversion of carbon dioxide and bicarbonate. CA inhibitors have been pursued for cancer, infectious disease, and ocular indications.^30–34^ For example, extracellular CA-II activity is problematic in diabetic retinopathy^35^ and CA-IX inhibition has exhibited preclinical and clinical efficacy in multiple cancers^36^. Numerous small molecule inhibitors demonstrate activity but modest specificity^33,34^, because of high isoform homology (Figure S1), which hinders fundamental study and causes detrimental side effects.^33^ A frequently studied small molecule, acetazolamide (AAZ), exhibits an inhibition constant (K_i_) in the tens of nM for several CA isoforms^33^ including 12 nM for CA-II and 25 nM for CA-IX. Extensive structural modifications have been explored to identify enhanced potency and/or improved selectivity, albeit with mixed results and substantial effort.^33^ A set of recently developed triazinyl-substituted benzenesulfonamides inhibit CA-II with ~10-fold selectivity (K_i_ ~ 13 nM) and CA-IX with ~700-fold selectivity (K_i_ ~ 0.15 nM), respectively^37,38^. These molecules were re-engineered to incorporate symmetric amino acid pairs into their tail regions and yield altered selectivity, producing a set of sulfonamides that are selective to CA-I, CA-IV, or CA-IX^39,40^. We integrate AAZ into a binding loop of the tenth type III domain of human fibronectin (Fn, also termed monobody or adnectin)^41^ as a cysteine-free antibody mimetic scaffold. Evolution of the amino acid sequence in its solvent-exposed loops has enabled development of specific, high affinity binders to a multitude of targets.^41,42^ Genetic incorporation of a single cysteine enables site-specific conjugation to maleimide pharmacophores.

The current study validates a platform for PriSM engineering via sorting a combinatorial library of 10^8^ Fn-AAZ hybrids to discover inhibitors of CA-II and CA-IX with superior potency and selectivity relative to small molecule or protein alone. Protein-small molecule synergy can be achieved by engineering various combinations of conjugation sites, linker lengths, protein sequences, and target isoforms. Beyond discovery of selective CA inhibitors, this ligand engineering platform provides broad opportunity for targeting epitopes to selectively elicit biological activity.

## Results

### PriSM library design and construction

We sought to engineer PriSMs with strong potency and isozyme selectivity to CA-II or CA-IX via modulation of the protein interface, AAZ conjugation site, and linker length. We designed combinatorial Fn libraries conserving a sole cysteine at a central, solvent-exposed site in either the central loop or the most protruding loop (sites 28 and 80 in the BC and FG loops, respectively; Figure 1A) and biased diversity within the adjacent three loops via guidance from previous protein evolution studies^43^. Eight sites were diversified with all 20 amino acids, biased toward tyrosine, serine, and other amino acids prevalent in antibody complementarity-determining regions and evolved Fn domains (Figure 1B, Figure S2); 11 sites had reduced diversity to balance inter- and intra-molecular constraints. Loop lengths were diversified to provide shape modulation: BC loop: −1, 0, +1 relative to wild-type; DE loop: −1, 0; FG loop: −2, −1, 0. Diverse genes, assembled from degenerate oligonucleotides, were homologously recombined into a *Saccharomyces cerevisiae* yeast display system as genetic fusions to N-terminal Aga2p with a semi-flexible polypeptide linker (Figure 1A). Both Cys28 and Cys80 libraries had 200 million unique transformants. Sequence analysis confirms that library construction generally matches design (median deviation from design (|*f*_observed_ – *f*_design_|) of 0.1% for both Cys28 and Cys80 libraries) (Figure S2).

**Figure 1.**
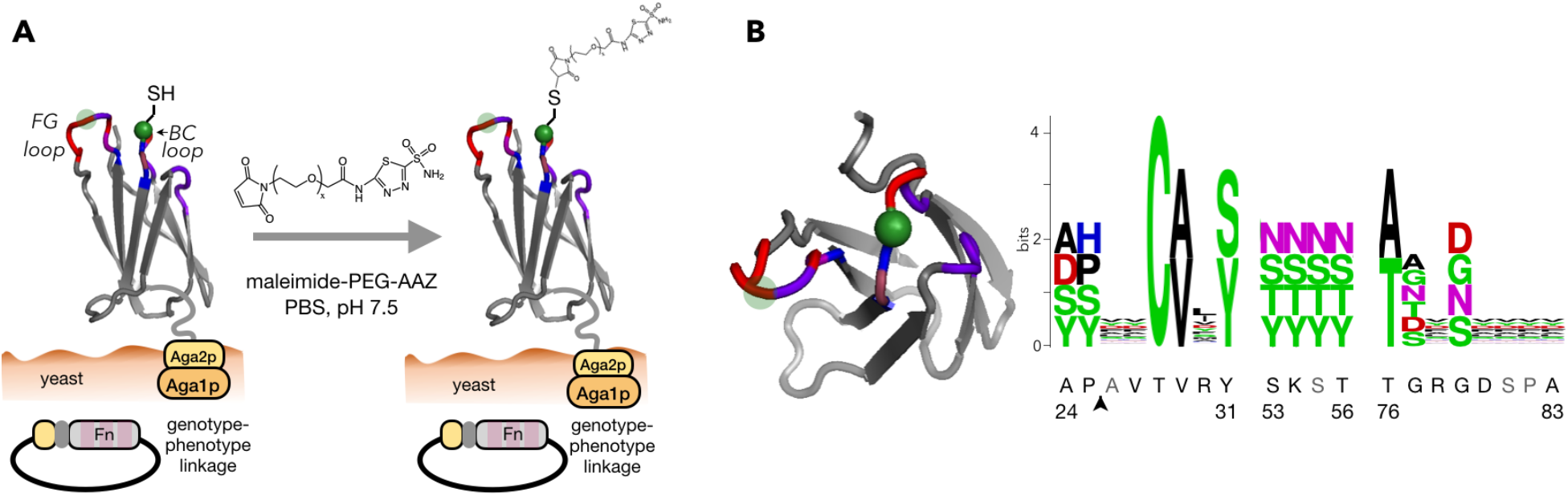
PriSM design integrates a pharmacophore with combinatorial protein diversity. (A) The Fn domain is genetically diversified at up to 19 sites in three solvent-exposed loops surrounding a conserved cysteine (green). Cys28 is shown explicitly with thiol; the semi-transparent green circle indicates site 80, which was alternatively conserved in a second library. Fn is tethered to the yeast cell wall via protein fusion to a semi-flexible peptide linker to Aga2p mating protein, which is bound to Aga1p anchored in the cell wall. Treatment with maleimide-PEG-AAZ yields the protein-small molecule conjugate PriSM. (B) *Left*: top view of Fn. *Right:* amino acid diversity at each site with wild-type residues indicated below (gray: deleted in shortened loop lengths; arrow: insertion in lengthened loops). The Cys28 library is shown; the Cys80 library is equivalent except the diversity at sites 28 and 80 are swapped.

### Engineered PriSMs exhibit potent, isoform-specific inhibition

Yeast were grown to replicate variants, induced to display Fn, and treated with maleimide-PEG_x_-AAZ (x = 2, 3, 5, or 7). We first describe the development of Fn-PEG-AAZ PriSMs that target CA-II and are conjugated at site 28 with a PEG_7_ linker. In later sections, we will show that this technology is also effective with an FG loop conjugation site, linker lengths between 2 and 7 PEG units, and targeting to inhibit CA-IX with isozyme specificity. The library was sorted twice for binding to biotinylated CA-II, and binders were pulled down with streptavidin-coated magnetic beads. Analysis of the sorted population revealed numerous binders of at least low affinity but also a majority of clones without detectable binding (Figure 2A). The presence of a multitude of non-binders highlights the need to precisely engineer the combination of protein sequence, conjugation site, linker, and pharmacophore for any target. Conversely, the presence of numerous binders indicates the potential for the PriSM concept to function in the context of a Fn protein scaffold, AAZ pharmacophore, and CA-II target.

**Figure 2.**
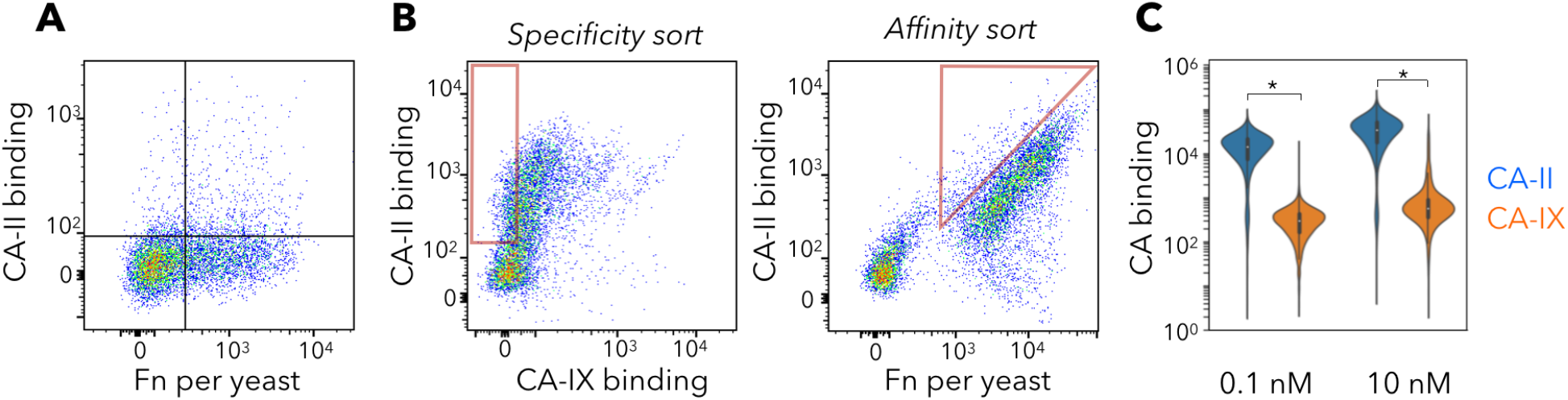
Selective, high-affinity PriSM discovery. (A) The magnetic bead-sorted population of yeast-displayed Fn was incubated with maleimide-PEG_7_-AAZ, washed, and incubated with 10 nM biotin-CA-II followed by streptavidin-AlexaFluor647. Fn display was quantified with mouse anti-Myc and anti-mouse-AlexaFluor488. Flow cytometric analysis of 10,000 random variants is shown. (B) The enriched population was labeled with DL650-CA-IX, biotin-CA-II+streptavidin-AF488, and mouse anti-Myc+anti-mouse-PE. Flow cytometric sorts enriched CA-II specific binders (left) with high affinity (right). (C) Final enriched population was labeled with CA-II or CA-IX at 0.1 or 10 nM each; binding was evaluated by flow cytometry. The distribution of 10,000 variants is presented. * indicates significant difference between binding CA-II and CA-IX (p < 0.001).

To promote increased diversity in subsequent sorts, Fn variants enriched from sorts with PEG_2_, PEG_3_, PEG_5_, and PEG_7_ versions of maleimide-PEG_x_-AAZ were pooled after each bead sort. The pooled population was sorted twice via stringent flow cytometric sorting – the first for affinity, the second for affinity and specificity. Yeast-displayed conjugates were incubated with both DyLight650-labeled CA-II and biotinylated CA-IX (detected by streptavidin-AF488). Conjugates that displayed high CA-II-DL650 labeling and low CA-IX-AF488 labeling were gated for sorting (Figure 2B). A second sort was performed with greater stringency to further enrich binders with high affinity and selectivity for CA-II. The resulting population was expressed on yeast, conjugated, and confirmed to selectively bind CA-II compared to CA-IX (Figure 2C).

Fn-encoding genes from the population enriched for strong, selective binders were shuttled to an *E. coli* expression vector. Clones were selected from this population at random, and Fn was expressed in the soluble fraction, purified via metal affinity chromatography, and conjugated with maleimide-PEG_7_-AAZ. Purity and conjugation were verified by SDS-PAGE and matrix-assisted laser desorption-ionization (MALDI) mass spectrometry (Figure S3).

Potency and specificity in inhibiting CA activity were evaluated via spectrophotometric monitoring of the conversion of 4-nitrophenyl acetate to 4-nitrophenol and acetic acid by carbonic anhydrase (Figure 3A).^44^ Fn_II.28.7_-PEG_7_-AAZ exhibits potent inhibition of CA-II (K_I_ = 4.8 nM) that is highly specific as CA-IX inhibition was 120-fold weaker (K_I_ = 580 nM; p = 0.01; Figure 3BC; confidence intervals are indicated in figures). The PriSM dramatically enhances specificity, while maintaining potency, compared to AAZ alone (K_I,CA-II_ = 3.3 nM; K_I,CA-IX_ = 6.6 nM). Moreover, the PEG linker impacts AAZ activity, thus it is also relevant to compare performance of the PriSM to unreacted PEG_7_-AAZ. The PriSM is 9-fold more potent (K_I,CA-II_ = 42 nM for PEG_7_-AAZ) and 80-fold more specific (K_I,CA-IX_ = 6 nM for PEG_7_-AAZ) than unreacted PEG_7_-AAZ. Notably, the evolved Fn_II.28.7_ protein alone, without PEG_7_-AAZ, does not inhibit CA-II activity. Thus, the PriSM concept enhances inhibitory performance in this context.

**Figure 3.**
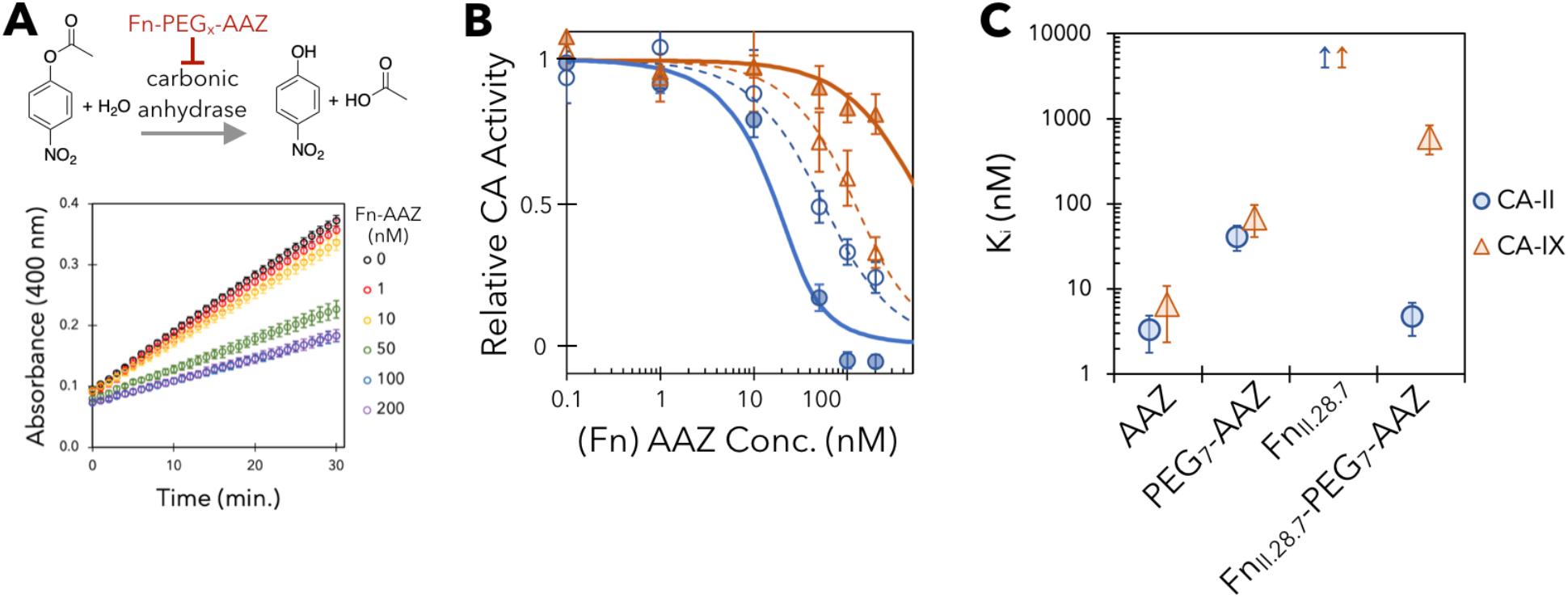
Engineered PriSMs exhibit potent, isoform-specific inhibition. (A) 4-nitrophenyl acetate is converted to 4-nitrophenol and acetic acid by carbonic anhydrase. The ability of PriSMs to inhibit this activity is titrated by spectrophotometrically measuring 4-nitrophenol production over time at different PriSM concentrations. (B) Titration curves of Fn_II-28.7_-PEG_7_-AAZ (solid lines) and PEG_7_-AAZ (dashed lines) with CA-II and CA-IX. (C) Inhibition constants. Data are presented as the mean ± 68% confidence interval for three replicates. Inhibition was not detected for Fn_II-28.7_.

### Potent, specific PriSMs can be engineered with multiple linker lengths, conjugation sites, and targets

To assess discovery of PriSMs with alternative linker length, analogous selection campaigns were performed with maleimide-PEG_x_-AAZ linkers of length x = 2, 3, and 5. Random colony picking of the enriched populations yielded clones with significant specificity for Fn_II.28.5_-PEG_5_-AAZ and Fn_II.28.3_-PEG_3_-AAZ, albeit not for the clone with PEG_2_ (Figure 4: *Cys28*). Fn_II.28.5_-PEG_5_-AAZ increases affinity 3.6-fold and specificity 9-fold relative to PEG_5_-AAZ. Fn_II.28.3_-PEG_3_-AAZ exhibits 11-fold specificity for CA-II relative to CA-IX by essentially eliminating observable inhibition of CA-IX but with an appreciable reduction in potency for CA-II. Thus, functional PriSMs can be engineered with a range of linker lengths. While it could be tempting to attribute reduced efficacy with shorter linker length, this is not the general case as shown by deeper analysis of linker length trends (see Figures 5C and S5) and demonstration of specificity and potency gains from short linkers in the context of other conjugation sites and isoform targets (Figure 4: *Cys80* and *CA-IX*, detailed next).

**Figure 4.**
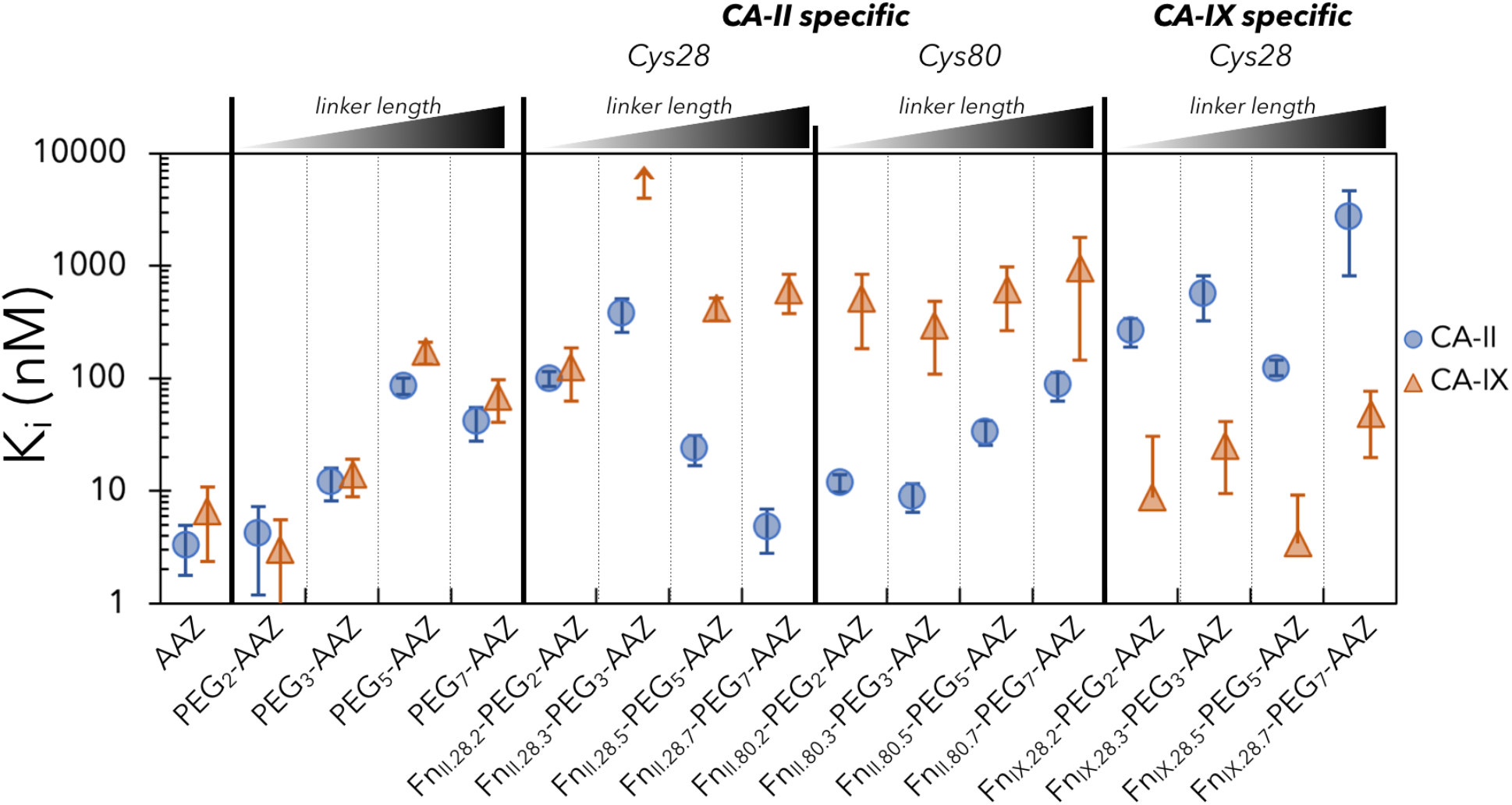
PriSMs can be engineered with multiple linker lengths. Enzymatic inhibition constants – measured via the assay detailed in Figure 3 – for AAZ, PEG-AAZ, and Fn-PEG-AAZ PriSMs. Clones were isolated from each of the four linker campaigns from the Cys28 and Cys80 libraries for CA-II binding and the Cys28 library for CA-IX binding. Data are presented as the mean ± 68% confidence interval for three replicates.

**Figure 5.**
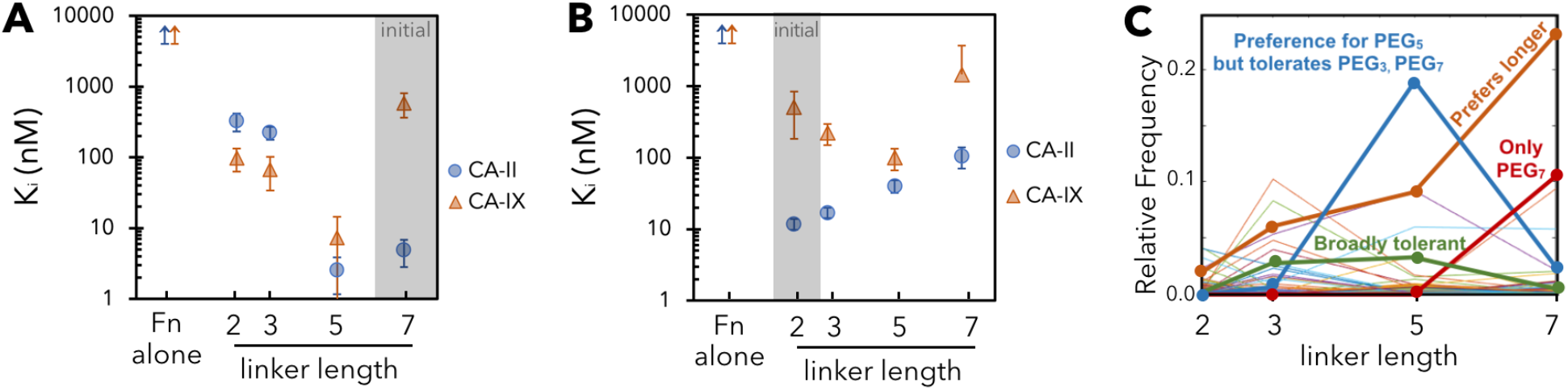
The PriSM-enzyme interface is finely tuned to linker length. (A,B) Modifying the linker from the length of original discovery hinders specificity. Enzyme inhibition constants for Fn_II.28.7_-L_x_-AAZ (A) and Fn_II.80.2_-L_x_-AAZ (B) (and both Fn proteins alone) are presented as the mean ± 68% confidence interval of triplicate measurements. Inhibition was not detected for Fn without conjugation. (C) Relative frequency across different linker length campaigns for CA-II binders from Cys80 library. Data exist at lengths 2, 3, 5, and 7; connecting lines are drawn for visual guidance.

To assess the ability to engineer PriSMs with an alternate protein conjugation site, binder selections were performed using the library conserving cysteine at site 80, conjugated to each of the four lengths of maleimide-PEG-AAZ, in parallel. Site 80 was chosen for its central location in the extended FG loop (Figure 1). A broad panel of specific, high-affinity binding variants was enriched (Figure S4). A selected clone at each linker length was evaluated for enzymatic inhibition. In all four cases, Fn conjugation engendered isoform specificity, with ratios of K_I_ values for CA-II to CA-IX of 11-42 (Figure 4: *Cys80*). Relative to PEG-AAZ alone, potency to CA-II was maintained (9 – 88 nM) while the engineered Fn drove the desired diminished activity against CA-IX. Thus, the Fn-PEG-AAZ concept is amenable to multiple Fn conjugation sites.

All conjugates to this point have been selected for specificity to CA-II compared to CA-IX. To assess the generality of the PriSM concept, we aimed to identify conjugates with the converse specificity. An analogous library selection scheme using magnetic bead and flow cytometric selections was used to isolate CA-IX binders. The selected conjugates achieved high potency (3-48 nM K_I_ values) and enhanced selectivity by 46-112 fold relative to small molecule alone (Figure 4: *CA-IX-specific*). Notably, Fn_IX.28.5_-PEG_5_-AAZ moderately enhanced potency to CA-IX relative to AAZ (to 3.4 nM from 6.6 nM) and drastically enhanced potency relative to PEG_5_-AAZ (170 nM), clearly demonstrating the addition of beneficial interactions from the Fn component. The performance of the collection of conjugates shows the applicability of the PriSM discovery platform to multiple targets.

### The PriSM-enzyme interface is finely tuned to linker length

Each aforementioned construct is associated with a particular PEG linker length with which it was discovered in the process of ligand selection. The intention of our design is that the AAZ pharmacophore inserts into the enzyme catalytic site, while the PEG linker creates space for the Fn component to find a low-energy, bound state. Specificity is the result of Fn finding energetically favorable interactions with its designed target, while generating unfavorable interactions with other non-targets. We hypothesize that a potent design constrains the linker configuration(s) to be neither too extended nor too collapsed (both low entropy states) while not hindering the binding interface. Notably, inclusion of the PEG linker, especially the longer two lengths tested, alters potency of AAZ (Figure 4). To determine whether the PEG linker length is an essential, constraining design parameter of each clone, we varied the linker length and assayed enzyme inhibition for Fn_II.28.7_-L_x_-AAZ and Fn_II.80.2_-L_x_-AAZ.

For Fn_II.28.7_-PEG_7_-AAZ, shortening the linker to 5 units results in a 40-fold specificity loss via dramatically elevating off-target CA-IX potency – either via removal of originally detrimental interactions or creation of beneficial interactions – while only nominally aiding CA-II potency (Figure 5A). Further reduction in linker length to 3 or 2 units drastically hinders CA-II potency by ~100-fold and also hinders CA-IX potency but less thereby resulting in a specificity reversal. The resulting potency toward both isoforms is substantially worse than the small molecule, with or without the linker, indicating that the Fn component, which was selected for enhancing potency with a longer linker, introduces detrimental interaction when constrained by a shorter linker. It is also notable, that the Fn protein alone does not provide measurable inhibition of either isoform.

Starting with a variant discovered to be effective with the shortest linker, Fn_II.80.2_-PEG_2_-AAZ, specificity reduction is again observed with linker modification. CA-II potency steadily worsens with longer linkers while CA-IX potency improves with extension to PEG_3_ and then PEG_5_ but is drastically hindered by further extension to PEG_7_ (Figure 5B). Also, as with the previous clone, the Fn protein alone is not inhibitory.

To broaden the analysis of linker length impact, we used the power of flow cytometry and deep sequencing. Each of the populations enriched during selection of high affinity, specific binders via flow cytometry (as in Figure 2B) were deep sequenced. The relative frequency of each variant reports on its efficacy; thus, comparison of frequencies across different length campaigns informs on linker length impact. An array of outcomes is observed (Figures 5C and S5). As a few examples from the CA-II binders with Cys80, one clone (orange in Figure 5C) performs better with longer PEG but tolerates all four lengths; another (blue) has substantially better performance with PEG_5_ but tolerates two units shorter or longer; another (red) is only effective with PEG_7_; conversely another (green) is broadly tolerant at all four lengths tested. Thus, use of only a single length is acceptable for efficiency as specific, high-affinity binders will be discovered. Yet a broader diversity of PriSM sequences will be discovered, including improvement of many clones, if length diversity is included in discovery.

### Analysis of the sequence landscape of functional PriSMs

Deep sequencing of selected populations also empowers analysis of the sequence landscape of functional PriSMs. Populations of engineered PriSMs are diverse, as evidenced by median Hamming distances of 9-14 within a campaign (Figure 6A). Most campaigns yielded moderate preferential enrichment of more functional variants with a consistent slope of 10-fold reduction in frequency per order of magnitude of diversity (Figure 6B). Notably, the clones randomly chosen from the final selection populations for inhibitory analysis were relatively broadly distributed. For example, the clones for PEG_5_ and PEG_7_ in Cys28 and PEG_7_ in Cys80 were the 3^rd^, 6^th^, and 4^th^ most frequent in their populations. Yet, other analyzed clones were 51^st^, 64^th^, 91^st^, and 240^th^. All were specific and potent except Fn^II.28.2^-PEG^2^-AAZ (which ranked 240^th^ thereby explaining its lack of efficacy on unfortunate stochastic selection). Given the specific potency of the other evaluated clones, there are clearly a broad array of functional PriSMs available.

**Figure 6.**
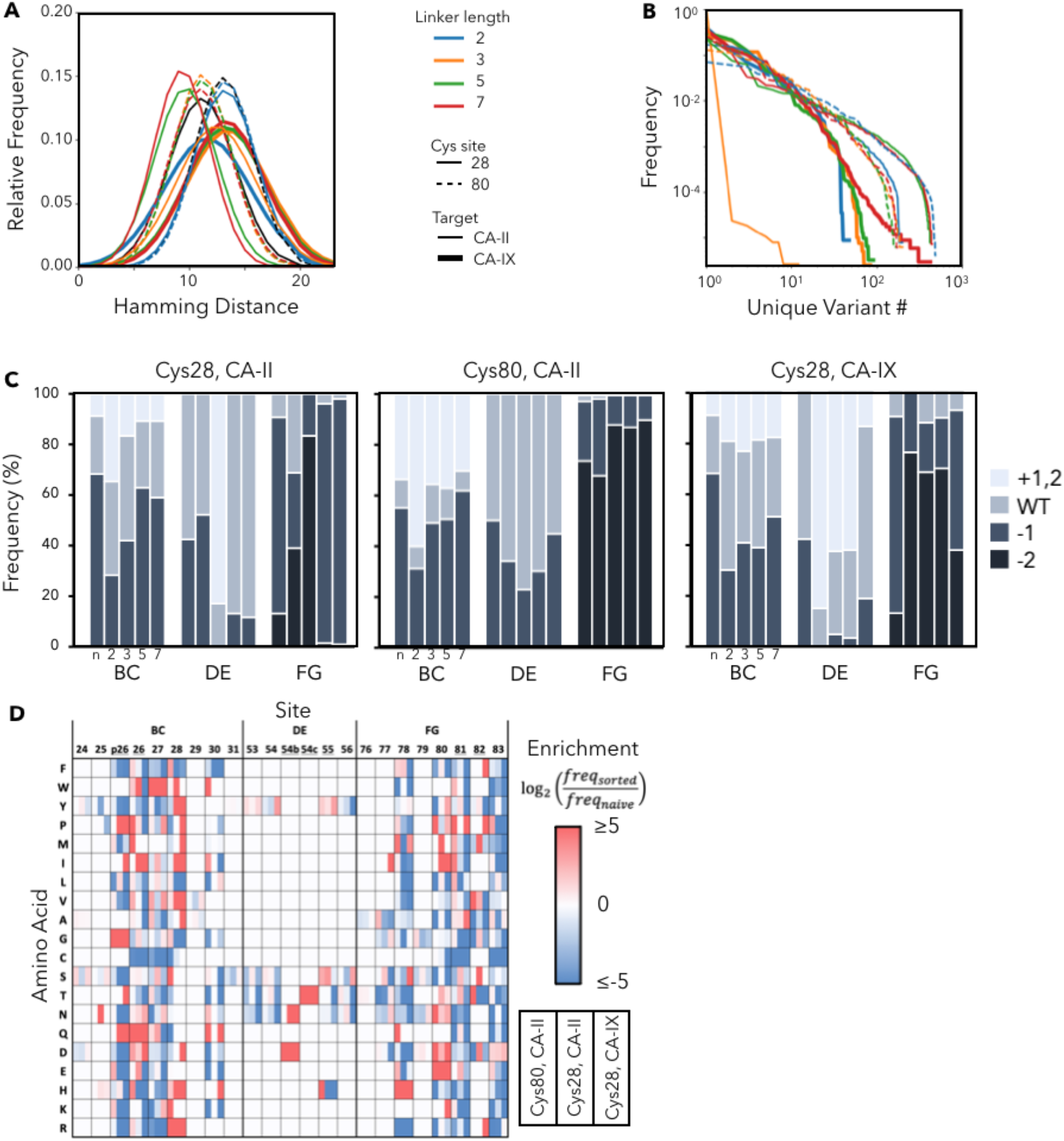
PriSM sequence landscape. (A) The Hamming distance between all pairs of variants in each population. Black represents the unsorted libraries. (B) The relative frequency of each variant in each population. (C) The relative frequency of each loop length for each campaign (Cys28 or Cys80 conjugation sites; naïve library or sorted for binding in the presence of conjugates with linkers of PEG_2_, PEG_3_, PEG_5_, or PEG_7_. (D) Sitewise amino acid enrichment, from the naïve library to the sorted binding population, is presented for each target (CA-II or CA-IX) and conjugation site (Cys28 or Cys80) combination, with results from all linker lengths aggregated. Sites with possible gaps for loop length diversity are underlined.

The functionally enriched populations were also evaluated for Fn loop length to guide future library design (Figure 6C). When containing the Cys28 conjugation site, BC loop length trends counter the PEG linker length, perhaps to provide sufficient cavity penetration for short linkers. The analog is not observed for FG loops with Cys80 conjugation, which is consistent with the already extended structure of the FG loop (Figure 1A). When AAZ is conjugated via short linkers to Cys28, short lengths are enriched in the adjacent FG loop, perhaps to avoid steric hindrance of the extended structure. Conversely, in Cys28 campaigns, longer DE loops are enriched, which could enable better target engagement of this otherwise smaller loop.

The PriSM sequence-function landscape is elucidated by analysis of amino acid enrichment at each site during selection of specific, high-affinity binders (Figure 6D). Cys is not enriched at other sites, which is consistent with the effective utility of the proposed conjugation sites C28 and C80. Multiple amino acid preferences at sites proximal to the AAZ conjugation site are revealed. Within Cys28 conjugation campaigns, site 27 is heavily enriched in W. Interestingly, this is counter to two other aromatics, the homologous F and Y, which are consistently depleted at the four sites flanking Cys28. At site 26, I, Q, and D are heavily enriched, which curiously represents a diverse chemical repertoire albeit with similar size; which may represent the importance of spatial packing at this location or diversity of engineered interfaces. Two sites from Cys28 AAZ conjugation on the other side, P30 is heavily enriched while numerous amino acids are substantially depleted. When the BC loop is at its longest, making p26 a non-gap amino acid, it is strongly enriched in G and P suggesting importance of loop conformation. In campaigns with Cys80 conjugation, the adjacent site 79 is enriched in G (from 17% to 39%), which may provide conformational flexibility. Conversely, the adjacent site on the other side, 81, is most heavily enriched in the conformationally constraining P. Lastly, the sites with initial diversity constraint (24, 25, 29, 31, 53-55, 76, 77, 79) generally do not exhibit substantial enrichment or depletion, which is consistent with effective library design; mild exceptions are enrichment of Y at sites 53 and 55 and S at 55. Collectively, analysis of the diverse PriSMs contributes to elucidation of sequence-function relationships that will be further enlightened by future campaigns and structural analysis.

## Discussion

The functional specificity and potency achieved herein highlight the power of hybridizing proteins and small molecule pharmacophores as well as the PriSM discovery platform. The systematic evaluation of hybrid performance across molecular design parameters demonstrates the generality of the approach as PriSM discovery was successful with two distinct conjugation sites in two different Fn loops, using four different PEG linker lengths, a broad diversity of protein sequences, and targeting two different enzyme isoforms. The distinctiveness of solutions across the different molecular designs reveals the impact of PriSM elements and underscores opportunities for further advancement via expansion and tuning of these design elements.

Notably, the high-throughput yeast display sorts were performed on the basis of target binding not inhibitory potency. Innovative platforms to efficiently sort for activity may aid discovery, yet activity-based selection with high throughput is a noted challenge of combinatorial discovery and directed evolution.^45,46^ A key strength of the PriSM concept, and the chemical biology discovery platform, may prove to be the ability of the pharmacophore to focus the search of molecular space on functionally active molecules (i.e. inhibitors in the current context, but could be agonists in other PriSM campaigns). Indeed, potent inhibition was achieved across numerous campaigns in the current work. Moreover, despite providing potency and selectivity enhancement, the engineered Fn domains without conjugated pharmacophore did not exhibit appreciable inhibition. The PriSM platform empowered efficient discovery of functional inhibitors. It is also striking that the current PriSMs were engineered without iterative evolution; that is, they were identified via direct discovery from a substantially undersampled set of sequence space (10^8^ experimentally tested genetic variants within a focused library design of 10^17^ possible protein variants with unbiased potential diversity of 10^26^).

In the 12 combinations of conjugation site, target, and linker length, specificity was always enhanced via hindrance of off-target potency. In 50% of these cases, on-target potency was also enhanced (Figure S6). The potency of the pharmacophore likely plays a role in this balance (reducing off-target potency versus enhancing on-target potency) when selecting for potent, selective PriSMs. Mutations more readily hinder interactions than aid^47^, thus a diverse population will tend to contain more variants that reduce off-target potency than enhance on-target potency. With a relatively strong pharmacophore, reduced off-target potency may be sufficient leading to enrichment of such variants. With a weaker pharmacophore, on-target potency enhancement becomes more requisite thereby increasing enrichment of such variants. Future efforts could explore the impact of pharmacophore affinity as well as other aforementioned design elements including linkers with different stiffness, hydrophilicity, or bulk; additional protein conjugation sites; and additional protein scaffolds. The yeast display cysteine-conjugation PriSM engineering platform demonstrated here can efficiently empower such studies.

## Significance

An armamentarium of diverse molecular architectures is needed to elicit precise biological function via active binding at select epitopes. While proteins, peptides, and small molecules have all exhibited efficacy in numerous cases, each format balances advantages and shortcomings. Previous work on peptide-pharmacophore hybrids motivates a complementary merger of formats to expand the molecular repertoire. Herein, we demonstrate the effectiveness of protein-small molecule hybrids or PriSMs as well as an effective platform for their discovery via yeast display library selections. The PriSM platform efficiently advances specificity and potency in selection from a single library without evolution. The success of the PriSM approach can be viewed from several lenses of drug discovery: a spatially integrated version of heterobivalent ligand design to boost avidity and specificity via dual binding interfaces; a hybrid approach to fragment-based drug discovery; or a synthetic chemistry approach to non-canonical amino acid incorporation. Demonstration of hybrid functionality with multiple linker lengths, conjugation sites, and protein sequences against a pair of isoforms highlights the potential to further tune molecular design from each of these perspectives. This study lays the foundation for future work to elucidate the fundamental understanding of these integrated interfaces as well as to apply the technological utility to a broad array of targets and biological functions.

## STAR Methods

### KEY RESOURCES TABLE

| REAGENT or RESOURCE | SOURCE | IDENTIFIER |
| --- | --- | --- |
| **Antibodies** |  |  |
| Mouse c-MYC Monoclonal Antibody (9E10) | Invitrogen | Cat# MA1-980, RRID AB_558470 |
| Streptavidin, Alexa Fluor™ 488 conjugate | Invitrogen | Cat# S11223 |
| Goat anti-Mouse IgG (H+L) Cross-Adsorbed Secondary antibody, PE | Invitrogen | Cat# P-852, RRID AB_2539848 |
| **Bacterial and Viral Strains** |  |  |
| *Escherichia coli*: DH5α | New England Biolabs | Cat# C2987C |
| *Escherichia coli*: T7 Express | New England Biolabs | Cat# C2566H |
| **Chemicals, Peptides, and Recombinant Proteins** |  |  |
| 4-Nitrophenyl acetate | Sigma-Aldrich | Cat# N8130-5G |
| DyLight™ 650 NHS Ester, 5 x 50 ug | Thermo Scientific | Cat# 62266 |
| EZ-Link™ Sulfo-NHS-LC-Biotin | Thermo Scientific | Cat# 21335 |
| HisPur™ Cobalt Resin | Thermo Scientific | Cat# 89964 |
| Human Carbonic Anhydrase II Protein (Recombinant) | LifeSpan BioSciences, Inc | Cat# LS-G551-100 |
| Human Carbonic Anhydrase IX / CA9 (38-414) Protein, His Tag | ACRO Biosystems | Cat# CA9-H5226-1mg |
| Carbonic Anhydrase IX Protein, Human, Recombinant (His Tag) | Sino Biological Inc. | Cat# 10107-H08H |
| Maleimide-PEG<sub>2</sub>-acetazolamide | Peptech | N/A |
| Maleimide-PEG<sub>3</sub>-acetazolamide | Peptech | N/A |
| Maleimide-PEG<sub>5</sub>-acetazolamide | Peptech | N/A |
| Maleimide-PEG<sub>7</sub>-acetazolamide | Peptech | N/A |
| **Critical commercial assays** |  |  |
| Zymoclean Gel DNA Recovery Kit | Zymo Research | Cat# D4002 |
| Zeba Spin Desalting column, 7 kDa | ThermoFisher | Cat# 89882 |
| LongLife Zymolyase | G-Biosciences | Cat#: 786-036 |
| **Experimental models: cell lines** |  |  |
| *Saccharomyces cerevisiae*: EBY100 | ATCC | MYA-4941 |
| **Oligonucleotides** |  |  |
| See Table S1 for PCR primers used in this study |  |  |
| **Recombinant DNA** |  |  |
| pCT40-FnLoopHP | Woldring et al., 2015 | N/A |
| pET-24b | Lipovšek et al., 2007 (Novagen) | N/A |
| **Software and Algorithms** |  |  |
| BioTek Gen5 Software Version 2.09 | Biotek | https://www.biotek.com/products/software-robotics/ |
| Matlab R2020a | MathWorks | https://www.mathworks.com/ |
| PyMol 2.4.0 Open-Source | PyMol | https://www.pymol.org/ |
| Python 3.8.8 | Python | https://www.python.org/ |

### RESOURCE AVAILABILITY

#### Lead Contact

Further information and requests for resources and reagents should be addressed to and will be fulfilled by the Lead Contact, Benjamin Hackel

#### Materials Availability

Key plasmids created in this study are available upon request.

#### Data and Code Availability

Raw source data are available upon reasonable request.

### EXPERIMENTAL MODEL AND SUBJECT DETAILS

#### Bacterial Strains

*Escherichia coli (E. coli)* DH5α strains were used for plasmid amplification. *E. coli* DH5α cells were grown in LB medium at 37 °C. *E. coli* T7 strains were used for protein purification for inhibition assays. Cells were grown at 37 °C until OD_600_ of 0.5-1.0, followed by induction with 0.5 mM isopropyl-β-d-thiogalactoside (IPTG) at 30 °C for 4 hours.

#### Cell Lines

*Saccharomyces cerevisiae* (*S. cerevisiae*) EBY100 cells were obtained from the American Type Culture Collection (ATCC). *S. cerevisiae* EBY100 cells were routinely cultured at 30 °C in rich YPD media. Once transformed with the yeast display pCT40-FnLoopHP plasmid (or library variants thereof), cells were grown in glucose-rich yeast minimal media minus tryptophan (SD-CAA) at 30 °C until OD_600_ of 6.0, followed by induction of yeast display with change into galactose-rich SG-CAA (0.1M sodium phosphate, pH 6.0, 6.7 g/L yeast nitrogen base, 5 g/L casamino acids, 19 g/L galactose, 1 g/L glucose).

### METHOD DETAILS

#### Materials and general methods

Synthetic DNA oligonucleotides (Table S1) were purchased from Integrated DNA Technologies (Coralville, IA). PCR and restriction digest products were purified by gel electrophoresis and extracted using Zymoclean Gel DNA Recovery kit. Plasmid DNA was purified using GenCatch Plasmid DNA Miniprep kit. Sanger DNA sequencing was conducted by Eurofins Genomics (Louisville, KY).

#### Library Construction

A genetic library was constructed based on the human tenth domain of fibronectin type III (Fn), in which the sequence encoding for the three loops was systematically diversified using degenerate oligonucleotides encoding for an amino acid distribution (Table S2). Degenerate nucleotides, at diversified positions 26-28, 78, and 80-83, had 20% A, 19% C, 27% G, and 34% T in the first position; 44% A, 21% C, 12% G, and 23% T in the second position; and 23% G, and 77% T in the third position as motivated by earlier library engineering^43^. The design also included loop length diversity with an insertion, deletion, or neither in the BC loop, a deletion or wild-type in the DE loop, and zero, one, or two deletions in the FG loop. Overlap extension PCR reactions were carried out to construct the full-length Fn genes. Gene reactions were transformed into EBY100 yeast using homologous recombination with linearized yeast surface display pCT40-FnLoopHP vector^43^.

#### Labeling of CA-II and CA-IX for FACS Detection

CA-II was first buffer exchanged with Zeba Spin Desalting Columns 7K MWCO to phosphate-buffered saline (PBS) and then incubated for four hours at 4 °C with 1:10 NHS-LC-biotin. CA-IX was reconstituted according to manufacturer recommendation and incubated for four hours at 4 °C with 1:100 with DyLight 650 NHS ester. Both conjugated CA-II and CA-IX were desalting by running twice through separate Zeba columns. Concentration was quantified via gel electrophoresis.

#### Maleimide-PEGx-AAZ on-yeast conjugation

Appropriate yeast populations were washed three times with PBS and then incubated for two hours at room temperature with 100 μM maleimide-PEG_x_-AAZ in 100 µL per 10 million cells. Conjugated yeast were then washed with PBSA (containing 0.1% BSA) to quench and clear any unbound maleimide-PEG_x_-AAZ.

#### Binder and specificity selection

Yeasts displaying the Fn Cys28 library were washed and conjugated in four campaigns to maleimide-PEG_x_-AAZ (x= 2, 3, 5, or 7) as described above. Conjugated yeast underwent magnetic activated cell sorting, where yeasts were incubated with control avidin-coated magnetic beads for one hour at room temperature, followed by another incubation of control avidin-coated beads to remove any non-specific binding interactions. Yeasts were then exposed to magnetic beads with immobilized CA-II biotinylated target protein and bound yeasts were selected. Magnetic sorts on the initial library were performed at room temperature and washed twice. Yeasts from each of the four linker length campaigns were pooled, grown, induced, and again conjugated, separately, to each of the four linker lengths. Non-naïve populations underwent a more stringent secondary magnetic sort. Conjugated yeasts were incubated twice with control avidin-coated beads, as before, incubated with 10 nM biotin-conjugated CA-II for 30 minutes, and then followed with incubation of avidin-coated beads for an additional 30 minutes for selection. Non-naïve populations were sorted at room temperature and washed four times. Two rounds of flow cytometry selections — the first with 10 nM biotinylated CA-II target, the second with 10 nM CA-II target and 10 nM DL650-labeled CA-IX non-target, and both with mouse anti-c-MYC and anti-mouse-PE — were used to isolate full-length (c-MYC positive) that bind selectively and with high affinity toward CA-II. The sort scheme was repeated for the Cys80 library.

Additionally, yeasts displaying the Fn Cys28 library were selected for CA-IX binders at each of the four PEG linker lengths. The naïve population first underwent the more stringent magnetic sort, instead incubated with 36 nM CA-IX biotinylated target, and selected with avidin-coated magnetic beads, as before. Yeasts were pooled, grown, induced, and reconjugated at each of the four lengths. Yeasts were sorted thrice via three-color FACS at 0.75 nM CA-IX and 50 nM CA-II, 0.5 nM CA-IX and 100 nM CA-IX, and again at 0.5 nM CA-IX and 100 nM CA-II.

#### Cloning, Protein Expression, and Purification

Fn-encoding regions in DNA recovered from the final CA-II/CA-IX flow cytometry sort were amplified by polymerase chain reaction (see Table S1), digested with NheI-HF and BamHI-HF reaction enzymes (New England Biolabs), and ligated with T4 DNA ligase into pET-24b vector2 containing a C-terminal hexa-histidine tag. Plasmids were transformed into T7 Express Competent *E. coli* and plated on lysogeny broth (LB) plates containing 50 mg/L kanamycin. Transformants were Sanger sequenced for full-length gene (Table S3), and proper transformants were grown in 5 mL liquid LB with kanamycin (50 mg/L) at 37 °C at 250 rpm for 12-16 hours. Saturated cultures were added to 100 mL LB, grown, and induced. Cells were pelleted and resuspended in lysis buffer (50 mM sodium phosphate (pH 8.0), 0.5 M sodium chloride, 5% glycerol, 5 mM 3-[(3-cholamidopropyl) dimethylammonio]-1-propanesulfonate, and 25 mM imidazole), frozen and thawed five times, centrifuged for 10 minutes at 4 °C, and 0.25 μm filtered. The resulting cell lysate were run through 0.25 mL Cobalt HisPur resin volume spin columns and conjugated to maleimide-PEG_x_-AAZ on-column (x = 2, 3, 5, or 7). The lysate-resin was washed with 30 mM imidazole, incubated with 1 mM TCEP for 15 minutes, washed with PBS, incubated for 30 minutes with 400 µM maleimide-PEG_x_-AAZ, washed with 30 imidazole, and then eluted with 300 mM imidazole. PriSMs were further purified via dialysis with three 1:1000 PBS buffer exchanges. Concentrations were quantified via NanoDrop One C (ThermoFisher Scientific, Madison, VA). PriSM purity was analyzed via gel electrophoresis, and small molecule conjugation was analyzed using AB-Sciex 5800 MALDI/TOF-MS.

#### Carbonic anhydrase inhibition assay

Carbonic anhydrase esterase activity was assayed by following the conversion of 4-nitrophenyl acetate (4-NPA) hydrolyzed to 4-nitrophenol and acetic acid in the presence of the enzyme via absorbance at 400 nm with a BioTek Synergy H1 microplate reader. CA-II or CA-IX was pre-incubated with inhibitor at various concentrations for 30 minutes prior to substrate addition, and the reaction was initiated upon addition of the substrate. The absorbance was measured every minute for 30 minutes at room temperature. The final concentration of the reaction components were 17.5 mM Tris, 105 mM NaCl buffer, pH 7.4, 25 nM CA-II or 100 nM CA-IX, 0, 1, 10, 50, 100, or 200 nM inhibitor, and 2 mM 4-NPA substrate in a final enzymatic assay volume of 100 µL containing 2% acetone. A higher concentration of CA-IX was used as CA-IX was less active than CA-II and therefore produced a much weaker signal. The absorbance of the 4-NPA with various inhibitor concentrations 0-200 nM in buffer, in the absence of enzyme, was subtracted from the enzymatic measurements. Each inhibitor was assayed with either CA-II or CA-IX with various concentrations at least three times.

#### Illumina MiSeq analysis of functional PriSMs

Plasmid DNA for the naïve Fn Cys28 and Fn Cys80 libraries as well as the final sorted populations was isolated from yeast using Zymolyase and extracted via silica spin column. The full Fn gene was amplified via PCR using full gene amplification primer (Table S1). Illumina adapters were added in a second PCR to differentiate between populations, and DNA was quantified by Nanodrop and mixed in equal molar ratios for a final concentration of 5 nM. Samples were submitted to the University of Minnesota Genomics Center for quality control analysis and sequenced using Illumina MiSeq (2×300, v3). Sequences were processed using USearch^48^ and filtered for a maximum of 1 total expected error for all bases in the read. Fn scaffold analysis was completed via homemade Python scripts to evaluate the frequency of unique variants, Hamming distance, loop length diversity, and site enrichment of final sorted populations.

### Quantification and Statistical Analysis

#### Determination of apparent KI

The absorbance values over time at different inhibitor concentrations were linearly fit to determine the slope. The fitted slope for various inhibitor concentrations in the absence of enzyme were used as a background subtraction. Inhibitor concentration did not affect substrate hydrolysis. Apparent K_i_ analyses were carried out in Matlab R2020a, fitting *v*_0_ and 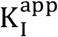 to Equation 1 using *nlinfit*. Default options were used for the solver except *TolX* and *TolFun* were both set to 1e-15. E_T_ was set as 25 nM for CA-II or 100 nM for CA-IX. Enzyme activity was then normalized to one using the fitted *v*_0_, the enzyme activity in the absence of inhibitor. The function *nlparci* was used to determine the 68% confidence interval for 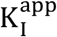, where alpha equaled 0.32.

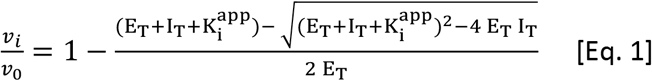

### Statistical analyses

Fluorescence single-cell data of final Fn Cys28-PEG_7_-AAZ population (Figure 2C) was exported, and a t-test was used to calculate p. Standard error of four independent experiments are shown in Fn_II.28.7_-PEG_7_-AAZ spectrophotometer data (Figure 3A) and titration curves (Figure 3B). P-values and standard error were calculated in Microsoft Excel.

## Supplemental Information

Figures S1-S6, Tables S1-S3

## Acknowledgements

We thank Alex Golinski for guidance on sequence analysis and Jim Van Deventer for useful discussions. We thank the University of Minnesota Flow Cytometry Resource for cell sorting, the University of Minnesota Genomics Center for assistance with deep sequencing, and the Minnesota Supercomputing Institute for computing resources. Research was supported by the National Institutes of Health (R01 EB023339) and Itara Biotherapeutics.

## Author Contributions

A.K.L., A.H., S.M.J., R.R.L., and B.J.H. designed research. A.K.L., A.H., and S.M.J. performed research. A.K.L., A.H., S.M.J., R.R.L., and B.J.H. analyzed data. A.K.L., A.H., and B.J.H. wrote the paper with input from all authors.

