## Supplemental Information for "Engineered protein-small molecule conjugates empower selective enzyme inhibition"

Figures S1-S6, Tables S1-S3

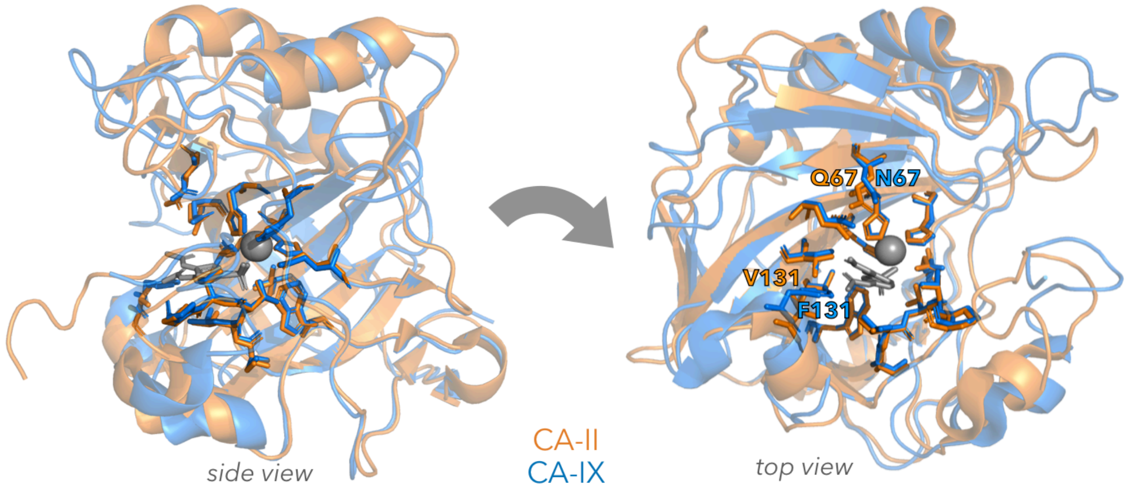


**Figure S1.** ***Carbonic anhydrase isoform homology.*** CA-II (blue) and CA-IX (orange) crystal structures with AAZ bound in the active site (PDB codes 3HS4 and 3IAI, respectively) were aligned. Side chains of amino acids within 5 Å of acetazolamide are shown and labeled.

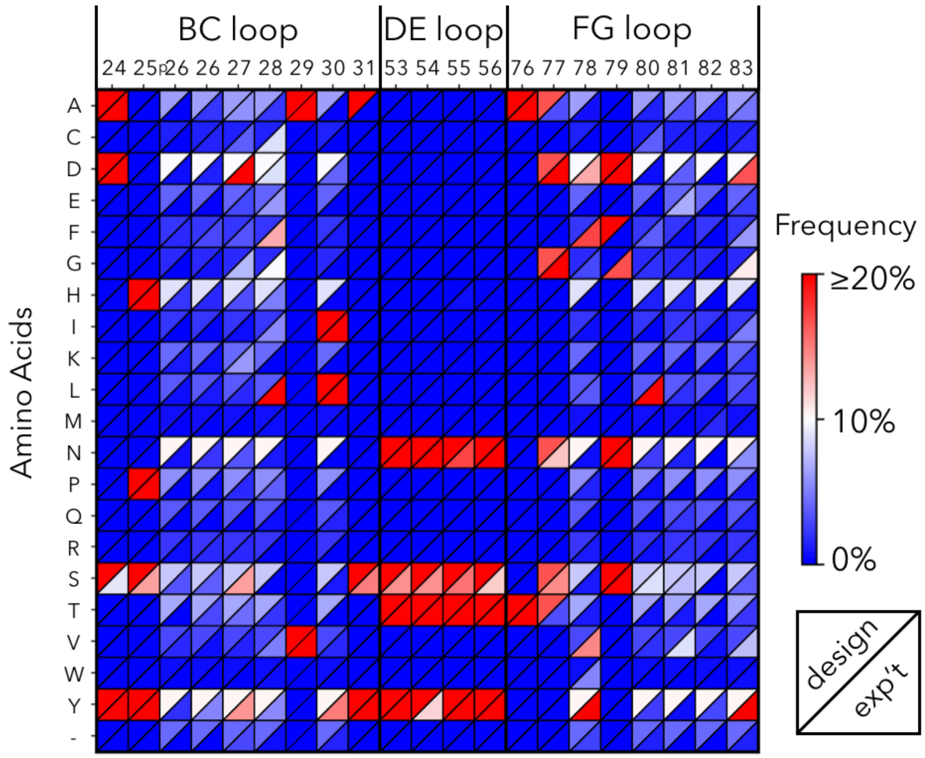


**Figure S2.** ***PriSM protein library matches design.*** Amino acid frequency at each site in the designed library (upper left triangle) and experimentally achieved library (lower right triangle). Data averaged across Cys28 and Cys80 libraries except sites 28 (only Cys80 diversity shown; 94% C in Cys28) and 80 (only Cys28 diversity shown; 89% C in Cys80). Median deviation is 0.1%.

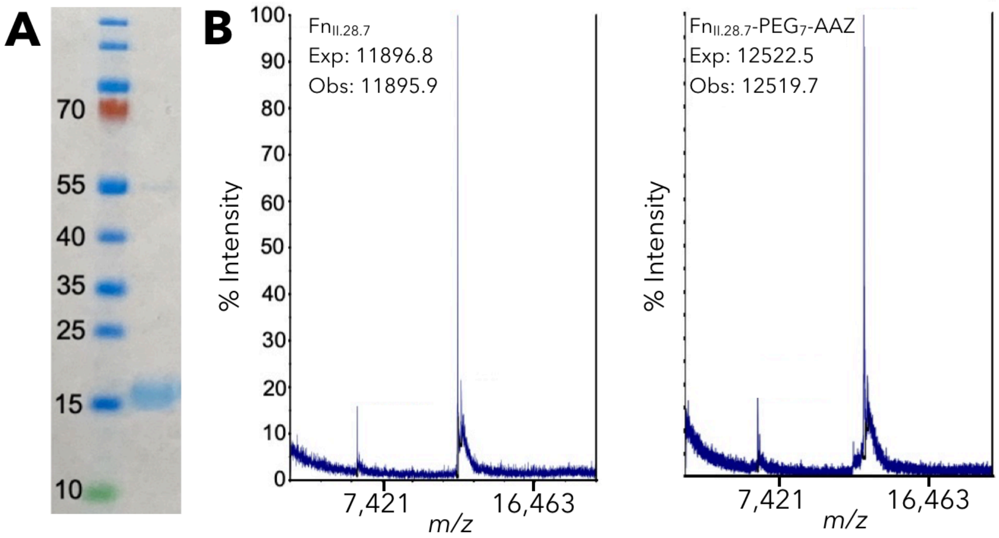


**Figure S3.** ***PriSMs are effectively produced via recombinant production and pharmacophore conjugation.*** (A) The Fn component of the $Fn_{II\text{-}28.7}$-$PEG_7$-AAZ PriSM was recombinantly produced in *E. coli*, purified by metal affinity chromatography, and analyzed via electrophoresis. (B) Conjugation to maleimide-$PEG_7$-AAZ is validated via matrix-assisted laser desorption ionization mass spectrometry.

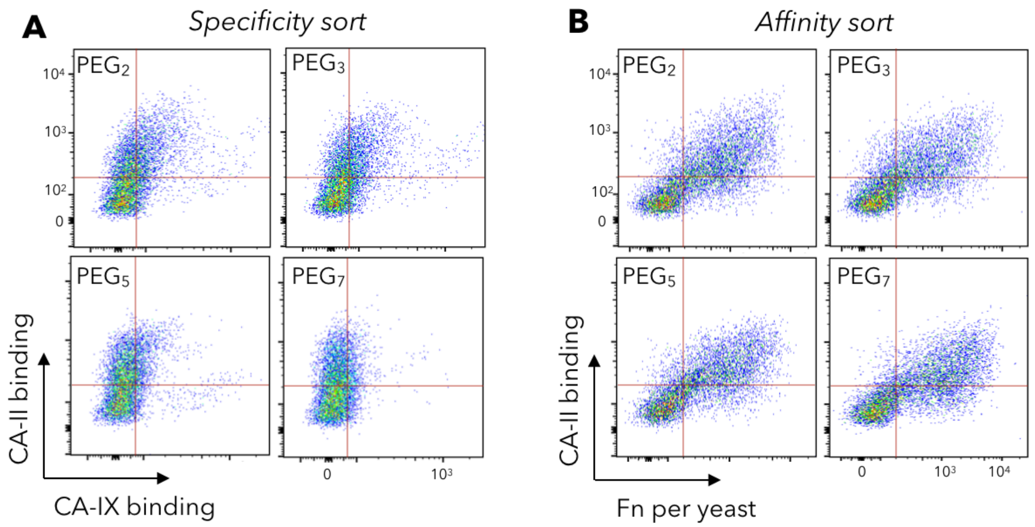


**Figure S4.** ***PEG-AAZ conjugation at Fn site 80 enables discovery of high affinity, isoform-specific PriSMs.*** The Cys80 Fn library was sorted for binding to CA-II analogously to the Cys28 library in Figure 2.

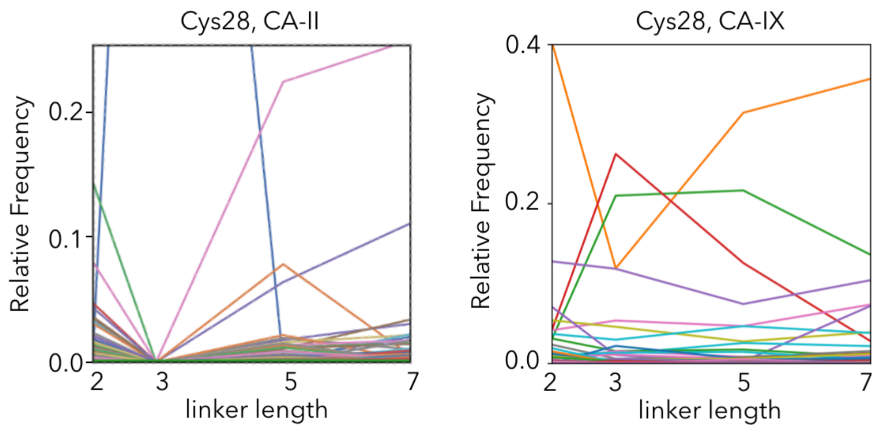


**Figure S5.** ***Clonal linker length preference.*** Relative clonal frequency across different linker length campaigns for CA-II and CA-IX binders from Cys28 library.

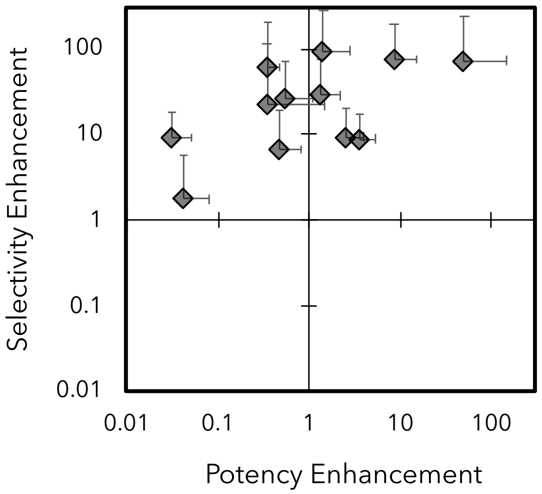


**Figure S6.** ***Conjugation to engineered Fn uniformly aids target selectivity and frequently aids target potency.*** The selected Fn-PEG-AAZ variant from each campaign (CA-II or CA-IX; Cys28 or Cys80; $PEG_2$, $PEG_3$, $PEG_5$, or $PEG_7$; Figure 4) is compared to $PEG_x$-AAZ for inhibitory potency ($K_i$). *Selectivity enhancement* is the ratio of on-target:off-target potency between Fn-PEG-AAZ and $PEG_x$-AAZ. *Potency enhancement* is the ratio of on-target potency between Fn-PEG-AAZ and $PEG_x$-AAZ.

**Table S1: DNA oligonucleotides for library construction and DNA amplification**

| Sequence | Description |
|---|---|
| ACTCTCTGACTATTTCTTGGGACKMTYMTxyzxyzxyzxyzGYTxyzTMTTACCGTATCACCTACGGCGAAAC | BC loop, length 10, 20* |
| ACTCTCTGACTATTTCTTGGGACKMTYMTxyzxyzxyzxyzGYTMTCTMTTACCGTATCACCTACGGCGAAAC | BC loop, length 10, IL |
| ACTCTCTGACTATTTCTTGGGACKMTYMTxyzxyzxyzGYTxyzTMTTACCGTATCACCTACGGCGAAAC | BC loop, length 9, 20* |
| ACTCTCTGACTATTTCTTGGGACKMTYMTxyzxyzxyzGYTMTCTMTTACCGTATCACCTACGGCGAAAC | BC loop, length 9, IL |
| ACTCTCTGACTATTTCTTGGGACKMTYMTxyzxyzGYTxyzTMTTACCGTATCACCTACGGCGAAAC | BC loop, length 8, 20* |
| ACTCTCTGACTATTTCTTGGGACKMTYMTxyzxyzGYTMTCTMTTACCGTATCACCTACGGCGAAAC | BC loop, length 8, IL |
| CGAGCCAGGAATTCACTGTTCCGGGAWMTWMTWMTWMTGCGACCATCAGCGGTCTGAAAC | DE loop, length 5 |
| CGAGCCAGGAATTCACTGTTCCGGGAWMTWMTWMTGCGACCATCAGCGGTCTGAAAC | DE loop, length 4 |
| CATTACCGTGTACGCTGTARCTRVTxyzRRCxyzxyzxyzxyzTCAAACCCAATCAGCATCAATTATCGCAC | FG loop, length 10 |
| CATTACCGTGTACGCTGTARCTRVTxyzRRCxyzxyzxyzTCAAACCCAATCAGCATCAATTATCGCAC | FG loop, length 9 |
| CATTACCGTGTACGCTGTARCTRVTxyzRRCxyzxyzTCAAACCCAATCAGCATCAATTATCGCAC | FG loop, length 8 |
| ACTCTCTGACTATTTCTTGGGACKMTYMTxyzxyzxyzTGCGYTxyzTMTTACCGTATCACCTACGGCGAAAC | BC loop, length 10, C28 w 20* |
| ACTCTCTGACTATTTCTTGGGACKMTYMTxyzxyzxyzTGCGYTMTCTMTTACCGTATCACCTACGGCGAAAC | BC loop, length 10, C28 w IL |
| ACTCTCTGACTATTTCTTGGGACKMTYMTxyzxyzTGCGYTxyzTMTTACCGTATCACCTACGGCGAAAC | BC loop, length 9, C28 w 20* |
| ACTCTCTGACTATTTCTTGGGACKMTYMTxyzxyzTGCGYTMTCTMTTACCGTATCACCTACGGCGAAAC | BC loop, length 10, C28 w IL |
| ACTCTCTGACTATTTCTTGGGACKMTYMTxyzTGCGYTxyzTMTTACCGTATCACCTACGGCGAAAC | BC loop, length 8, C28 w 20* |
| ACTCTCTGACTATTTCTTGGGACKMTYMTxyzTGCGYTMTCTMTTACCGTATCACCTACGGCGAAAC | BC loop, length 8, C28 w IL |
| CATTACCGTGTACGCTGTARCTRVTxyzRRCTGCxyzxyzxyzTCAAACCCAATCAGCATCAATTATCGCAC | FG loop, length 10, C80 |
| CATTACCGTGTACGCTGTARCTRVTxyzRRCTGCxyzxyzTCAAACCCAATCAGCATCAATTATCGCAC | FG loop, length 9, C80 |
| CATTACCGTGTACGCTGTARCTRVTxyzRRCTGCxyzTCAAACCCAATCAGCATCAATTATCGCAC | FG loop, length 8, C80 |
| CGACGATTGAAGGTAGATACCCATACGACGTTCCAGACTACGCTCTGCAG | Vector |
| ATCTCGAGCTATTACAAGTCCTCTTCAGAAATAAGCTTTTGTTCGGATCC | Vector |
| CGACGATTGAAGGTAGATACCCATACG | Full gene amplification primer |
| ATCTCGAGCTATTACAAGTCCTCTTC | Full gene amplification primer |

* xyz = CDR diversity (x = 20% A, 19% C, 27% G, 34% T)(y = 44% A, 21% C, 12% G, 23% T)(z = 00% A, 00% C, 23% G, 77% T);

**Table S2:** *Library design.* The wild-type Fn sequence and library diversity is indicated at each diversified loop position. 20* refers to a biased composition that balances amino acid frequencies observed in human antibody complementary-determining regions, as well as evolved Fn domains. Loop lengths were varied by including or excluding codons at sites pre-26, 26, 55, 81, or 82 as indicated by -. Cysteine is conserved at either site 28 or site 80. At site 30, an equal mixture of two codons, 20* and IL, was used.

| | BC Loop | | | | | | | | | DE Loop | | | | FG Loop | | | | | | | |
|---|---|---|---|---|---|---|---|---|---|---|---|---|---|---|---|---|---|---|---|---|---|
| Name | 24 | 25 | p26 | 26 | 27 | 28 | 29 | 30 | 31 | 53 | 54 | 55 | 56 | 76 | 77 | 78 | 79 | 80 | 81 | 82 | 83 |
| Wild-type | A | P | - | A | V | T | V | R | Y | S | K | S | T | T | G | R | G | D | S | P | A |
| Library (Cys28) | ADSY | HPSY | 20*/- | 20*/- | 20* | C | AV | 20*/IL | SY | NSTY | NSTY | NSTY/- | NSTY | AT | ADGNST | 20* | DGNS | 20* | 20*/- | 20*/- | 20* |
| Library (Cys80) | ADSY | HPSY | 20*/- | 20*/- | 20* | 20*/IL | AV | 20*/IL | SY | NSTY | NSTY | NSTY/- | NSTY | AT | ADGNST | 20* | DGNS | C | 20*/- | 20*/- | 20* |

**Table S3: Amino acid sequences for engineered Fn binders**

| Variant | Amino Acid Sequence |
|---|---|
| $Fn_{II.28.2}$ | MASSSDSPRNLEVTNATPNSLTISWDDPKCAYYYRITYGETGGNSPSQEFTVPGNTNATISGLKPGQDYTITVYAVTGFSSLSNPISINYRTEIDKPSQGSHHHHHH |
| $Fn_{II.28.3}$ | MASSSDSPRNLEVTNATPNSLTISWPWICPGMYHMPISINYRTEIDKPSQGSHHHHHH |
| $Fn_{II.28.5}$ | MASSSDSPRNLEVTNATPNSLTISWDYPDCAIYYRITYGETGGNSPSQEFTVPGTNTNATISGLKPGQDYTITVYAVTDYSLSNSNPISINYRTEIDKPSQGSHHHHHH |
| $Fn_{II.28.7}$ | MASSSDSPRNLEVTNATPNSLTISWDDSAACALYYRITYGETGGNSPSQEFTVPGTSYNATISGLKPGQDYTITVYAVADYSLQYSNPISINYRTEIDKPSQGSHHHHHH |
| $Fn_{II.80.2}$ | MASSSDSPRNLEVTNATPNSLTISWDDPAFAYYYRITYGETGGNSPSQEFTVPGYTNNATISGLKPGQDYTITVYAVTAEDCDSNPISINYRTEIDKPSQGSHHHHHH |
| $Fn_{II.80.3}$ | MASSSDSPRNLEVTNATPNSLTISWDYSPYGAYYYRITYGETGGNSPSQEFTVPGYNSTATISGLKPGQDYTITVYAVTDFGCDSNPISINYRTEIDKPSQGSHHHHHH |
| $Fn_{II.28.5}$ | MASSSDSPRNLEVTNATPNSLTISWDASGFAYYYRITYGETGGNSPSQEFTVPGYSSYATISGLKPGQDYTITVYAVTDFNCFSNPISINYRTEIDKPSQGSHHHHHH |
| $Fn_{II.80.7}$ | MASSSDSPRNLEVTNATPNSLTISWDYHYLALYYRITYGETGGNSPSQEFTVPGNTYATISGLKPGQDYTITVYAVTDYDCISNPISINYRTEIDKPSQGSHHHHHH |
| $Fn_{IX.28.2}$ | MASSSDSPRNLEVTNATPNSLTISWDYHQNGCAVSYRITYGETGGNSPSQEFTVPGYYDTYSATISGLKPGQDYTITVYAVTGYNDDSNPISINYRTEIDKPSQGSHHHHHH |
| $Fn_{IX.28.3}$ | MASSSDSPRNLEVTNATPNSLTISWDDHDKCVIYYRITYGETGGNSPSQEFTVPGYYDTYSATISGLKPGQDYTITVYAVANSDEDSNPISINYRTEIDKPSQGSHHHHHH |
| $Fn_{IX.28.5}$ | MASSSDSPRNLEVTNATPNSLTISWDDYQNCAVSYRITYGETGGNSPSQEFTVPGYYDTYSATISGLKPGQDYTITVYAVTGYNDDSNPISINYRTEIDKPSQGSHHHHHH |
| $Fn_{IX.28.7}$ | MASSSDSPRNLEVTNATPNSLTISWDDYLLCVFSYRITYGETGGNSPSQEFTVPGYTNTVTISGLKPGQDYTITVYAVTTYDLDSNPISINYRTEIDKPSQGSHHHHHH |
